# Prediction of Biopharmaceutical Characteristics of PROTACs using the ANDROMEDA by Prosilico Software

**DOI:** 10.1101/2022.09.22.509053

**Authors:** Urban Fagerholm, Sven Hellberg, Jonathan Alvarsson, Ola Spjuth

**Affiliations:** Prosilico AB, Lännavägen 7, SE-141 45 Huddinge, Sweden; Department of Pharmaceutical Biosciences and Science for Life Laboratory, Uppsala University, Box 591, SE-751 24 Uppsala, Sweden

## Abstract

**Background:** PROTACs are comparably large and flexible compounds with limited solubility (S) and permeability (P_e_). It is crucial to better understand, predict and optimize their human clinical pharmacokinetics (PK).

**Methods:** The main objective was to use the ANDROMEDA by Prosilico software to predict the human clinical *in vivo* dissolution potential (f_diss_) and fraction absorbed (f_a_) of 23 PROTACs at a dose level of 50 mg and to explore whether there is any relationship between *in vitro* S and *in silico* predicted *in vivo* f_diss_.

**Results:** *In silico* predictions showed that the PROTACs are effluxed by intestinal transporters and have limited f_diss_ (34 to 98 %), permeability and f_a_ (13 to 58 %) in man. For some PROTACs this may be a major obstacle and jeopardize the clinical development programs, especially in cases of required high oral dose. A modest relationship between *in vitro* S and predicted *in vivo* f_diss_ was demonstrated (R^2^=0.26). Predicted human f_a_ (27 %) and oral bioavailability (20 %) of ARV-110 (a PROTAC with some available *in vivo* PK data in rodents and man) were consistent with data obtained in rodents (estimated f_a_ approximately 30-40 %; measured oral bioavailability 27-38 %). Laboratories were unable to quantify S for 7 (30 %) of the PROTACs. In contrast, ANDROMEDA could predict parameters for all.

**Conclusion:** ANDROMEDA predicted f_diss_ and f_a_ for all the chosen PROTACs and showed limited f_diss_, P_e_ and f_a_ and dose-dependent f_diss_ and f_a_. One available example shows promise for the applicability of ANDROMEDA for predicting biopharmaceutics of PROTACs *in vivo* in man. Weak to modest correlations between S and f_diss_ and a considerable portion of compounds with non-quantifiable S limit the use of S-data to predict the uptake of PROTACs.

## Introduction

PROTACs are heterobifunctional molecules built of three moieties (a warhead binding a protein of interest, an E3 ligase recruiter and a linker attaching both regions) characterized by comparably large and flexible structures (1,2). Low solubility (S) and permeability (P_e_) are assumed to hinder their gastrointestinal uptake and systemic exposure following oral administration.

Jiménez et al. (1) measured thermodynamic S of 21 PROTACs and built *in silico* methodology for prediction, classification and optimization of S. Their selected PROTACs have S ranging between <0.23 and 661 µM. Their suggested and used cut-off values for low, intermediate and high S are <30 (n=13), 30-200 (n=5) and >200 (n=3) µM, respectively (1).

A limitation with the study by Jiménez et al. is that dose and the interplay between S, P_e_ and dose were not considered (this needs to be taken into account when predicting oral uptake). Thomas et al. (3) proposed minimum S-limits of ca 20, 200 and 2000 µM for minimum acceptable uptake of low P_e_-compounds in man at oral doses of 0.1, 1 and 10 mg/kg (ca 7, 70 and 700 mg), respectively (3).

Another limitation with the study by Jiménez et al. is the apparently low clinical relevance of *in vitro* S for *in vivo* S and dissolution. Poor correlations between S in phosphate buffer and human intestinal fluid (R^2^<0.2 (4)) and between S and maximum *in vivo* dissolution potential (f_diss_; the fraction of oral dose dissolved in the human gastrointestinal tract during optimized conditions) (R^2^=0.01 (data on file)) have been shown, and there is also extensive interlaboratory variability for S (2.3- and 100-fold median and mean variability, respectively (5)).

Our prediction software ANDROMEDA by Prosilico predicts human clinical PK, including parameters such as f_diss_, fraction absorbed (f_a_), oral bioavailability (F) and efflux by MDR-1, BCRP and MRP2. ANDROMEDA, with a major domain for molecules with molecular weight (MW) of 100-700 g/mole, is based on conformal prediction (CP), which is a methodology that sits on top of machine learning methods and produce valid levels of confidence (6), unique algorithms and a new human physiologically-based pharmacokinetic (PBPK) model (7). For a more extensive introduction to CP we refer to Alvarsson et al. 2014 (8). ANDROMEDA has been validated in several studies and shown to outperform laboratory methods in accuracy and breadth (9-14). The software has been applied to predict the f_diss_ and f_a_ of macrocyclic compounds with MW above its major domain (MW up to 1203 g/mole) (12). A 2-fold median prediction error for f_a_ was reached. There was, however, a slight underprediction trend, which may have been due to factors such as compound size above the upper MW-threshold for the software, impact of molecular flexibility on absorption at higher MW, uncertainties in measured estimates, saturation effects and contribution by active uptake. Demonstrated advantages with ANDROMEDA include ability to predict absorption/biopharmaceutics for macrocycles with low P_e_ and recovery in the Caco-2 cell model and very low solubility.

In line with the macrocyclic compounds, PROTACs have MW near or above 700 g/mole (699 to 1179 g/mol in the study by Jiménez et al (1)).

Apparently, human clinical uptake data for PROTACs are lacking, and this limits the opportunity to validate prediction methods. There are, however, PK-data in rodents for PROTAC ARV-110 that strongly indicate that limited gastrointestinal S and P_e_ are main reason behind the low F and that this is also expected in humans. The oral F of ARV-110 in rats and mice has been estimated to 27 and 38 %, respectively (15). A half-life of 11-18 h in rodents (15) and 110 h in man (16) indicates good metabolic stability *in vivo*.

The main objective of the study was to use ANDROMEDA to predict the human clinical biopharmaceutics (f_diss_ and f_a_) of ARV-110 and ARV-471 and the 21 PROTACs selected by Jiménez et al. at an oral dose level of 50 mg and to explore whether there is any relationship between *in vitro* S and *in silico* predicted *in vivo* f_diss_.

## Methods

### Compounds and solubility data

The 23 PROTACs (including *in vitro* thermodynamic S measured with shake-flask method at pH 7 and 25 °C and reported by Jiménez et al. (1), MW and SMILES) selected for the study are shown in Tables 1 and 2. Their S (available for all compounds except for ACBI1, ARV-110, ARV-471, ARV-825, CisACBI1, Mcl1 degrader-1 and MD-224 due to quantification problems) and MW range from <0.23 to 661 µM and 699 to 1179 g/mole (average 937 g/mole).

**Table 1.**
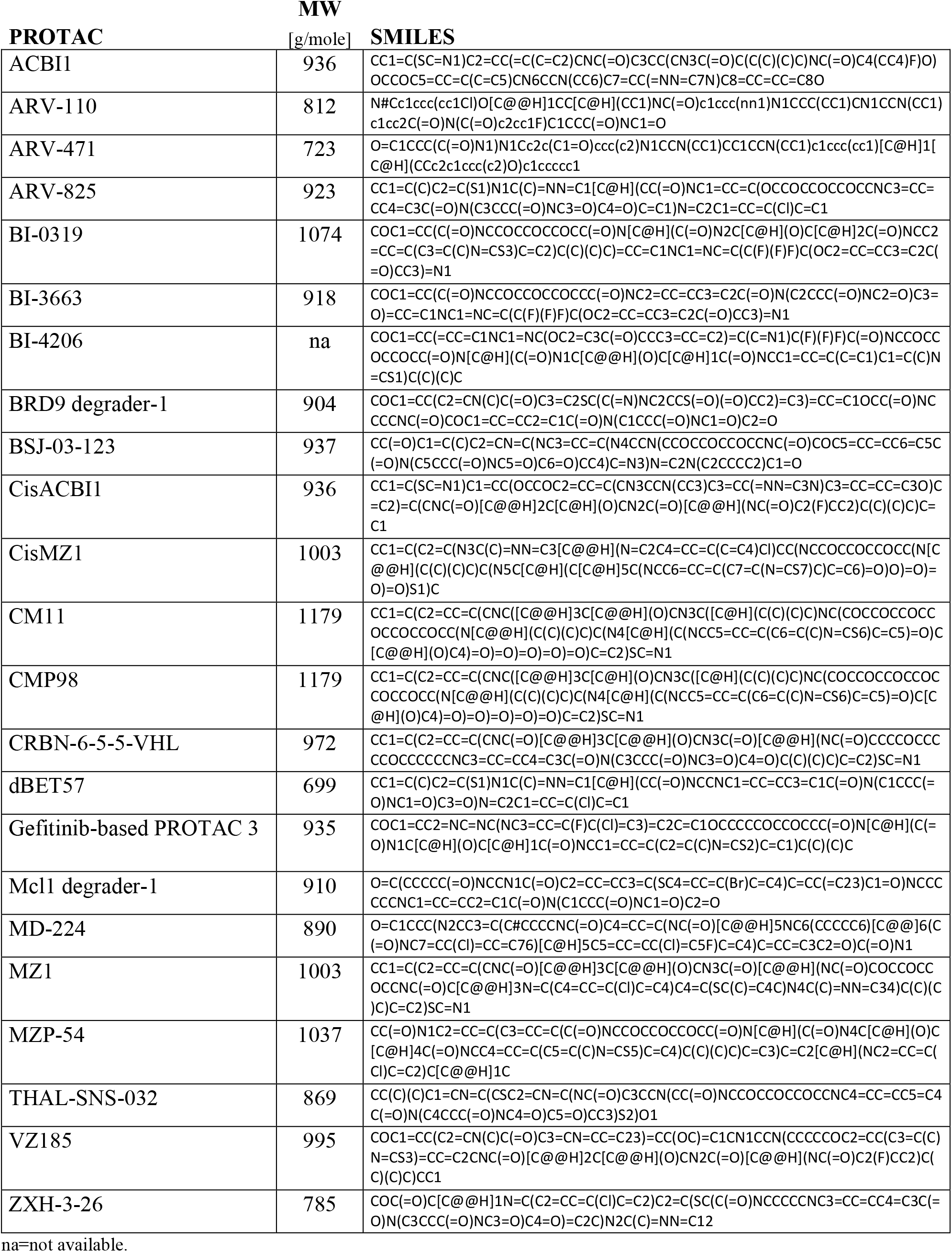
Selected PROTACs and their molecular weights and SMILES.

**Table 2.** Selected PROTACs and their *in vitro* S (solubility) and *in silico* predicted *in vivo* f_diss_ (*in vivo* dissolution potential) at an oral dose level of 50 mg.

| PROTAC | S<br>[μmol/L] | f<sub>diss</sub>*<br>[fraction] |
| --- | --- | --- |
| ACBI1 | <0,72 | 0,98 |
| ARV-110 | na | 0,65 |
| ARV-471 | na | 0,62 |
| ARV-825 | <0,32 | 0,45 |
| BI-0319 | 2,6 | 0,66 |
| BI-3663 | 6,9 | 0,44 |
| BI-4206 | 0,58 | 0,72 |
| BRD9 degrader-1 | 660,7 | 0,64 |
| BSJ-03-123 | 17,8 | 0,58 |
| CisACBI1 | <0,87 | 0,98 |
| CisMZ1 | 50,1 | 0,84 |
| CM11 | 631,0 | 0,84 |
| CMP98 | 331,1 | 0,84 |
| CRBN-6-5-5-VHL | 1,2 | 0,42 |
| dBET57 | 30,2 | 0,71 |
| Gefitinib-based PROTAC 3 | 1,2 | 0,63 |
| Mcl1 degrader-1 | <0,87 | 0,34 |
| MD-224 | <0,23 | 0,39 |
| MZ1 | 38,0 | 0,84 |
| MZP-54 | 0,51 | 0,68 |
| THAL-SNS-032 | 52,5 | 0,71 |
| VZ185 | 50,1 | 0,70 |
| ZXH-3-26 | 3,0 | 0,56 |

na=not available. \*at an oral dose level of 50 mg.

### Parameters predicted with ANDROMEDA

The prediction system based on CP and a new PBPK-model (ANDROMEDA software by Prosilico for prediction, simulation and optimization of human clinical PK) was applied to predict the main parameters for the study, f_a_ and f_diss_ (at 50 mg oral dose). Q^2^ values (cross-validation results for predicted *vs* observed data) obtained for passive f_a_, f_diss_ and F with *in silico* methodology developed by our team are 0.8, 0.6 and 0.5, respectively (7). Substance-relevant f_a_ and f_diss_ based on predicted or actual clinical dose can be calculated using an additional algorithm. Additional parameters were MDR-1, BCRP and MRP2 substrate specificity, f_a_ by passive permeation and oral F. None of the selected compounds in the present study was included in the training sets. Thus, every prediction was a forward-looking prediction where each compound was unknown to the software.

## Results

Prediction results are shown in Table 2 and Figures 1 and 2. According to the *in silico* predictions, all the PROTACs are effluxed by MDR-1, BCRP and/or MRP2, their f_diss_ and f_a_ at 50 mg oral doses level range between 34 and 98 % (average 66 %) and 13 and 58 % (average 34 %), respectively.

**Figure 1.**
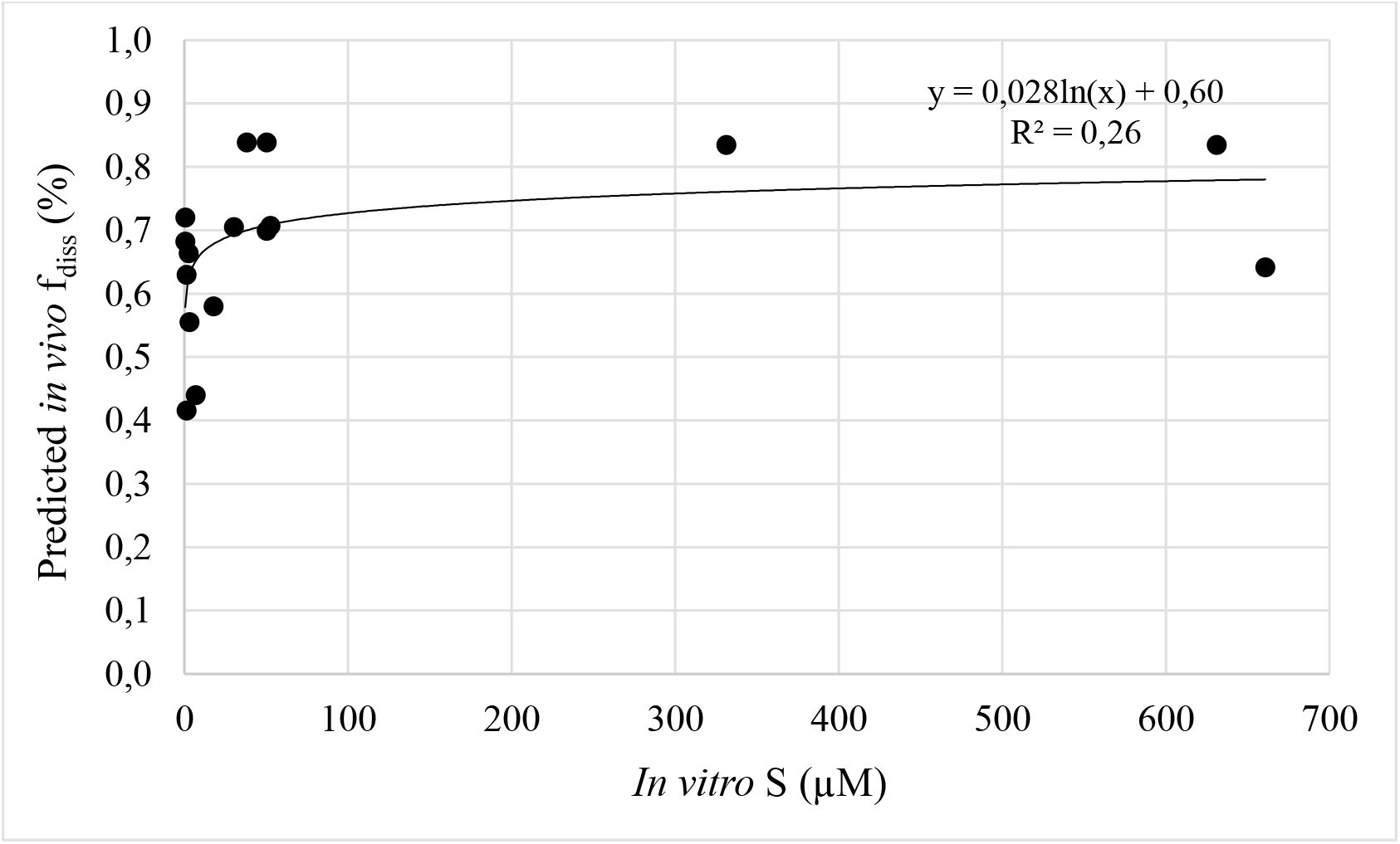
The relationship between *in vitro* S and *in silico* predicted *in vivo* f_diss_ (n=16).S for ACBI1, cisACBI1, ARV-825, Mcl1 degrader-1 and MD-224 were below the quantification limit (<0.32-0.87 µM) and S-estimates for ARV-110 and ARV-471 were unavailable.

**Figure 2.**
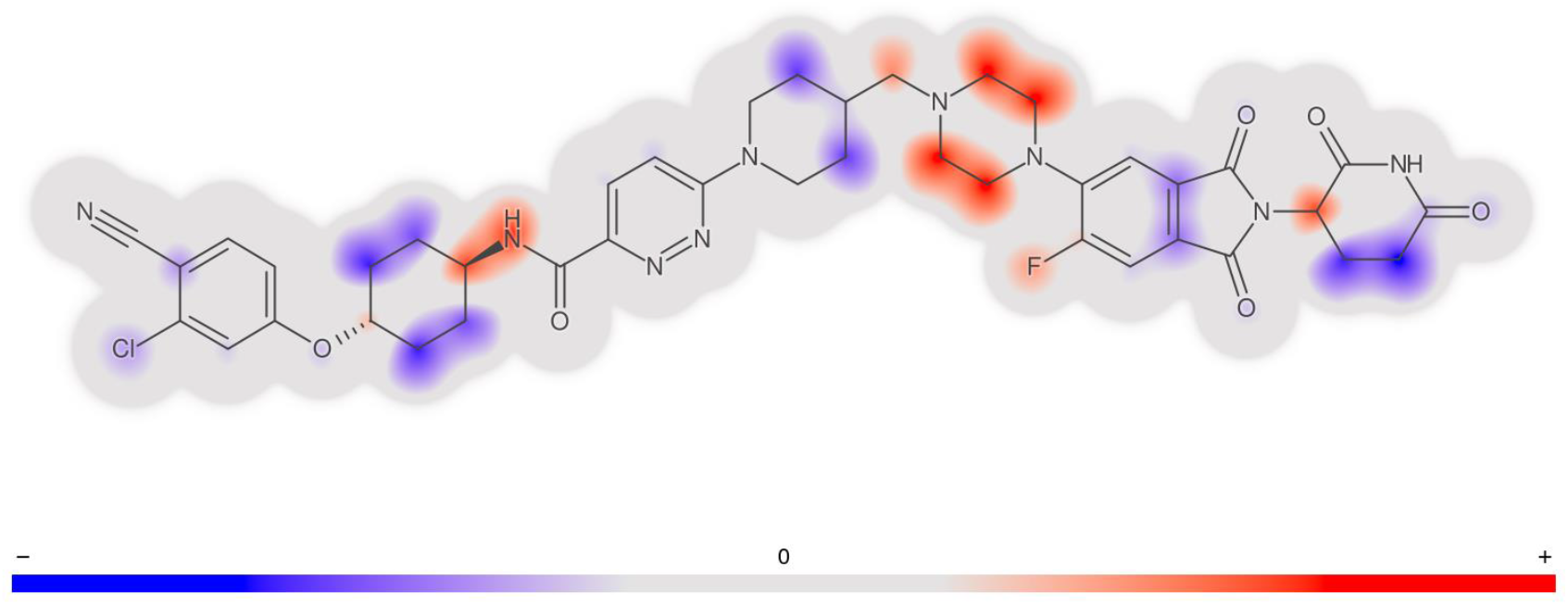
Molecular structure and signatures for ARV-110. Molecular regions contributing to decreasing (blue) and increasing (red) the f_diss_ are shown.

Lowest f_diss_ was predicted for Mcl1 degrader-1 (34 %), which has an *in vitro* S and MW of <0.87 µM and 910 g/mole. The corresponding estimates for the compound with highest f_diss_, ACBI1, are 98 % (100 % at 1 mg dose), <0.72 µM and 936 g/mole, respectively.

The predicted f_diss_ for the 3 compounds with the highest *in vitro* S, BRD9 degrader-1, CM11 and CMP98, were 64, 84 and 84 %, respectively.

Passive P_e_-based f_a_ (excluding efflux and solubility/dissolution) ranged between 50 and 88 %.

For ARV-110, f_diss_, f_a_ and F at 50 mg oral dose were predicted to 65, 27 and 20 %, respectively. Predicted f_diss_ at 1 and 500 mg oral doses were ca 90 and 50 %, respectively.

Nine of the 23 compounds had a predicted F<20% and the lowest individual F was 3 %.

There was an apparent relationship (R^2^=0.26) between *in vitro* S and *in silico* predicted *in vivo* f_diss_.

## Discussion

Prediction results show that PROTACs are effluxed by MDR-1, BCRP and/or MRP2 and have limited f_diss_ (at 50 mg oral dose), P_e_, f_a_ and F in man (BCS class 2 or 4 depending on oral dose size). For some PROTACs this may be a major obstacle and jeopardize clinical development programs, especially in cases of required high oral dose.

A modest relationship between *in vitro* S and *in silico* predicted *in vivo* f_diss_ was demonstrated (R^2^=0.26), indicating that *in silico* predicted f_diss_ estimates can be used to make rough predictions of *in vitro* S obtained with the specific methodology and that S-data alone are weak predictors of *in vivo* f_diss_. A poorer S-f_diss_ relationship may be found when using S-data from a different or a mixture of methods. For example, there was virtually no correlation between S and f_diss_ for a larger set of small molecules with low to high S and f_diss_ (data on file). R^2^-values at oral dose levels 1 and 500 mg were lower than at 1 mg, which demonstrates the impact and importance of dose for predictions of *in vivo* solubility and dissolution.

There is very little human clinical uptake and PK-data for PROTACs available, which makes it difficult to validate prediction methods. Data in rodents (oral F of 27-38 % and good metabolic stability) indicate that the f_a_ of ARV-110 is approximately 30-40 %, which is similar to predicted values in man (27 % f_a_ and 20 % F).

The consistency between predictions in man and measurements in rodents for ARV-110 indicates potential of ANDROMEDA for predicting biopharmaceutical profiles for PROTACs. The ability of ANDROMEDA to predict *in vivo* f_diss_ at different dose levels and with 70 % confidence intervals (12)) and to guide towards optimized PK-properties is attractive. The choice of signature descriptors as machine learning features allows for highlighted molecular structural features that contribute most to predictions, which is a decision aid in optimization of PK properties (see Figure 2 where molecular structure and signatures for the f_diss_ of ARV-110 are demonstrated). In addition, the software is useful when laboratories fail to measure and quantify certain characteristics, such as for S for 7 (30 %) of the 23 PROTACs in this study.

ANDROMEDA can be further evaluated when/if more clinical biopharmaceutical/PK data for PROTACs become available.

## Conclusion

ANDROMEDA predicted f_diss_ and f_a_ for all the chosen PROTACs and showed limited f_diss_, P_e_ and f_a_ and dose-dependent f_diss_ and f_a_. One available example shows promise for the applicability of ANDROMEDA for predicting and optimizing biopharmaceutics of PROTACs *in vivo* in man. Weak to modest correlations between S and f_diss_ and a considerable portion of compounds with non-quantifiable S limit the use of S-data to predict the uptake of PROTACs.

